# Cell cycle-specific loading of condensin I is regulated by the N-terminal tail of its kleisin subunit

**DOI:** 10.1101/2022.08.19.504508

**Authors:** Shoji Tane, Keishi Shintomi, Kazuhisa Kinoshita, Yuko Tsubota, Tomoko Nishiyama, Tatsuya Hirano

## Abstract

Condensin I is a pentameric protein complex that plays an essential role in mitotic chromosome assembly in eukaryotic cells. Although it has been shown that condensin I loading is mitosis-specific, it remains poorly understood how the robust cell cycle regulation of condensin I is achieved. Here we set up a panel of *in vitro* assays to demonstrate that cell cycle-specific loading of condensin I is regulated by the N-terminal tail (N-tail) of its kleisin subunit CAP-H. Deletion of the N-tail accelerates condensin I loading and chromosome assembly in *Xenopus* egg mitotic extracts. Phosphorylation-deficient and phosphorylation-mimetic mutations in the CAP-H N-tail decelerate and accelerate condensin I loading, respectively. Remarkably, deletion of the N-tail enables condensin I to assemble mitotic chromosome-like structures even in interphase extracts. Together with other extract-free functional assays *in vitro*, our results uncover one of the multilayered mechanisms that ensure cell cycle-specific loading of condensin I onto chromosomes.

## Introduction

Chromosome assembly is an essential cellular process that ensures equal segregation of the genetic information into two daughter cells during mitosis (Batty & Gerlich, 2019; Paulson *et al*, 2021). Extensive studies during the past two decades or so have established the consensus that the condensin complexes play a central role in this process (Hirano, 2016; Uhlmann, 2016). Many if not all eukaryotic species have two different condensin complexes, known as condensins I and II, each of which is composed of five subunits. The two condensin complexes share a pair of SMC (structural maintenance of chromosome) ATPase subunits (SMC2 and SMC4) but are distinguished by distinct sets of non-SMC subunits (Ono *et al*, 2003). In condensin I, the kleisin subunit CAP-H bridges the head domains of a V-shaped SMC2-SMC4 heterodimer, and is then bound by a pair of HEAT subunits, CAP-D2 and -G (Fig 1A, right). In vertebrates, the two condensin complexes cooperate to assemble rod-shaped chromosomes in mitosis (Ono *et al*, 2003; Shintomi & Hirano, 2011; Green *et al*, 2012; Gibcus *et al*, 2018), but they display differential subcellular localization during the cell cycle. In HeLa cells, for instance, condensin I primarily resides in the cytoplasm during interphase, and gets loaded onto chromosomes immediately after the nuclear envelope breaks down in prometaphase (Ono *et al*, 2004; Hirota *et al*, 2004). However, mitosis-specific loading of condensin I can be recapitulated in membrane-free *Xenopus* egg extracts (Hirano *et al*, 1997), suggesting that a mechanism(s) independent of the regulation of subcellular localization also operates to ensure the mitosis-specific action of condensin I. Thus, a whole molecular picture of how the loading and action of condensin I are tightly regulated throughout the cell cycle remains to be determined.

**Figure 1.**
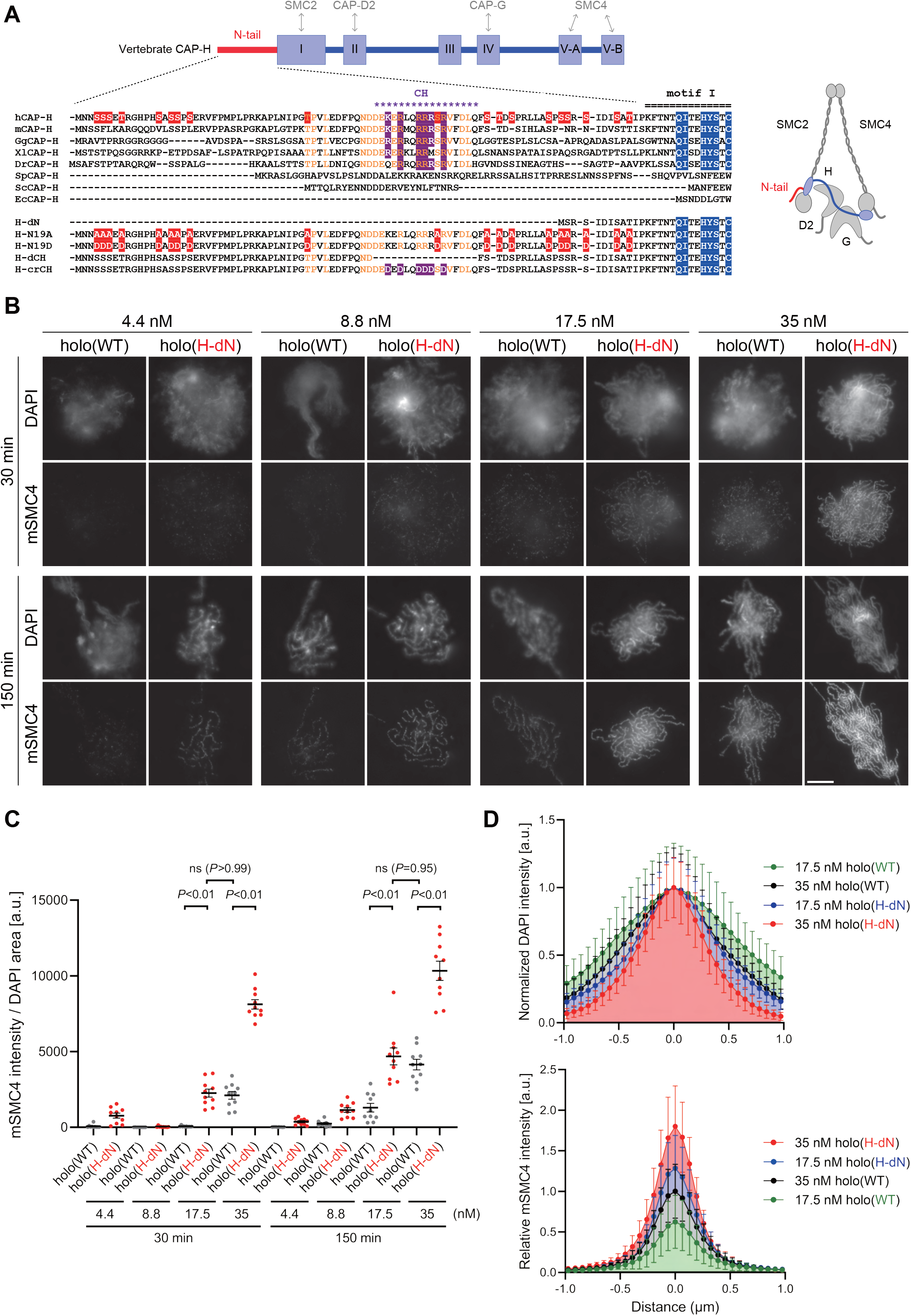
Deletion of the CAP-H N-tail accelerates condensin I loading and mitotic chromosome assembly. (A) Domain organization of vertebrate CAP-H and sequence alignments of the CAP-H N-tail in eukaryotes (left). Shown in the upper half is an alignment of the CAP-H orthologs (*Homo sapiens:* hCAP-H, *Mus musculus:* mCAP-H, *Gallus gallus:* GgCAP-H, *Xenopus laevis:* XlCAP-H, *Danio rerio:* DrCAP-H, *Schizosaccharomyces pombe:* SpCAP-H, *Saccharomyces cerevisiae:* ScCAP-H, and *Encephalitozoon cuniculi:* EcCAP-H). Six mutants tested in the current study (H-dN, H-N19A, H-N19D, H-dCH, and H-crCH) are shown in the bottom half. Conserved amino-acid residues were shown in yellow (N-tail) or blue (motif I). The helix motif predicted using Jpred4 (http://www.compbio.dundee.ac.uk/jpred/) is indicated by “CH” (for the conserved helix). The N19A/N19D and crCH mutation sites were shown in red and purple, respectively. Also shown in a schematic diagram of the architecture of vertebrate condensin I (right). (B) Add-back assay with holo(WT) and holo(H-dN). Mouse sperm nuclei were incubated with condensin-depleted M-HSS that had been supplemented with holo(WT) or holo(H-dN) at concentrations of 4.4, 8.8, 17.5, and 35 nM. After 30 and 150 min, the reaction mixtures were fixed and processed for immunofluorescence labeling with an antibody against mSMC4. DNA was counterstained with DAPI. Scale bar, 10 μm. (C) Quantification of the intensity of mSMC4 per DNA area in the experiment shown in Fig 1B (n = 10 clusters of chromosomes). The error bars represent the mean ± SEM. The *P* values were assessed by Tukey’s multiple comparison test after obtaining a significant difference with two-way ANOVA. (D) Line profiles of mitotic chromosomes observed at 150 min shown in the experiment shown in Fig 1B. Signal intensities of DAPI (top) and mSMC4 (bottom) from chromosomes assembled by holo(WT) or holo(H-dN) at 17.5 or 35 nM were measured along with the lines drawn perpendicular to chromosome axes (n = 20). The mean and standard deviation were normalized individually to the DAPI intensities (arbitrary unit [a.u.]) at the center of chromosome axes (distance = 0 μm)(top). Intensities of mSMC4 signals were normalized relative to the value from holo(WT) at 35 nM (bottom). The error bars represent the mean ± SD.

Early studies demonstrated that condensin I is phosphorylated in a mitosis-specific manner in *Xenopus* egg extracts (Hirano *et al*, 1997), and that the positive supercoiling activity of condensin I can be activated by Cdk1 phosphorylation *in vitro* (Kimura *et al*, 1998; Kimura *et al*, 2001). More recently, a mitotic chromatid reconstitution assay using purified proteins was used to demonstrate that Cdk1 phosphorylation of condensin I allows the complex to load onto chromosomes and to drive chromatid assembly (Shintomi *et al*, 2015).Phosphorylation of condensin I by other mitotic kinases has also been reported (Lipp *et al*, 2007; Takemoto *et al*, 2007; St-Pierre *et al*, 2009). All subunits of condensin I are large polypeptides of >100 kD, making it a daunting challenge to identify all phosphorylation sites and dissect their functional impacts. A global proteomic analysis of Cdk1 phosphorylation sites provided a clue to potentially relieving this problem, however (Holt *et al*, 2009).This study showed that Cdk1 phosphorylation often occurs in intrinsically disordered regions (IDRs) of proteins whose evolutionary conservation is relatively poor.It seems that this rule can be applied to the subunits of condensin I as well (Bazile *et al*, 2010).

In the current study, we focus on the N-terminal IDR of the vertebrate kleisin subunit CAP-H. We show that this region, which is referred to as the CAP-H N-tail, acts as a negative regulatory element for condensin I function. Deletion of the N-tail accelerates condensin I loading and mitotic chromosome assembly in *Xenopus* egg extracts. Phosphorylation-deficient mutations in the N-tail decelerate condensin I loading, whereas phosphorylation-mimetic mutations accelerate this process. Remarkably, when the N-tail function is compromised, the resultant mutant forms of condensin I enable the assembly of mitotic chromosome-like structures even in interphase extracts. The N-tail deletion mutant is also characterized in topological loading and loop extrusion assays *in vitro*. Taken together, our results uncover one of the multilayered mechanisms of cell cycle regulation of condensin I.

## Results and Discussion

### The N-terminal tail of the kleisin subunit CAP-H

The kleisin subunit CAP-H of condensin I has five sequence motifs widely conserved among eukaryotes (Fig 1A) (Hara *et al*, 2019; Kinoshita *et al*, 2022). Among them, motif I and V bind to the SMC neck and the SMC4 cap regions, respectively (Hassler *et al*, 2019), whereas motif II and motif IV interact with CAP-D2 and CAP-G, respectively (Piazza *et al*, 2014; Kschonsak *et al*, 2017; Hara *et al*, 2019). In the current study, we focus on the N-terminal extension, located upstream of motif I, that is shared by the vertebrate CAP-H orthologs (Fig 1A). This extension, the CAP-H N-tail, has several characteristic features as summarized below. Firstly, it is ~80 amino-acid-long in vertebrates, but the corresponding extension is much shorter or completely missing in fungi (Bazile *et al*, 2010). Secondly, although most of the N-tail is predicted to be structurally disordered, a stretch of ~17-amino-acid-long, located in its middle region, is conserved among vertebrates and is predicted to form an α-helix, as judged by both conventional secondary structure prediction software Jpred4 (https://www.compbio.dundee.ac.uk/jpred/) and AlphaFold2 (https://alphafold.ebi.ac.uk/). Thirdly, the vertebrate N-tail contains multiple SP/TP sites, sites often targeted by cyclin-CDK complexes, although their number varies among different species.

### Deletion of the CAP-H N-tail accelerates condensin I loading and mitotic chromosome assembly

To address the role of the CAP-H N-tail in condensin I function, we expressed mammalian subunits of condensin I using a baculovirus expression system in insect cells, and purified recombinant complexes according to the procedure described previously (Kinoshita *et al*, 2015; Kinoshita *et al*, 2022). In the first set of experiments, a holocomplex containing wild-type hCAP-H, holo(WT), and a holocomplex containing mutant hCAP-H that lacks its N-terminal 77 amino acids, holo(H-dN), were prepared (Fig 1A and EV1A). We then tested the ability of these recombinant complexes to assemble mitotic chromosomes in *Xenopus* egg extracts (high-speed supernatants of metaphase-arrested extracts, hereafter referred to as M-HSS) that had been immunodepleted of endogenous condensin subunits (Kinoshita *et al*, 2015; Kinoshita *et al*, 2022) (Fig EV1C). The condensin-depleted extracts were supplemented with increasing concentrations (4.4 nM, 8.8 nM, 17.5 nM, and 35 nM) of holo(WT) or holo(H-dN), mixed with mouse sperm nuclei and incubated at 22°C. Aliquots were taken from the reaction mixtures at 30 min and 150 min, fixed and processed for immunofluorescence using an antibody against the recombinant subunit mSMC4 (Fig 1B). We found that holo(WT) produced a cluster of rod-shaped chromosomes with mSMC4-positive axes under the standard condition (i.e., 35 nM condensin I, 150-min incubation) set up in the previous study (Kinoshita *et al*, 2022). As expected, progressively poorer assembly was observed with decreasing concentrations of condensin I (17.5-4.4 nM) or at the earlier time point (30 min). Remarkably, we found that significantly higher levels of holo(H-dN) were detected on chromatin than holo(WT) at both time points and at the different concentrations of condensin I tested (Fig 1B and C). It was also noticed that the chromosome sassembled with 35 nM holo(H-dN) at 150 min were much thinner than those assembled with 35 nM holo(WT) at 150 min (Fig 1B and Fig 1d). When the concentration of holo(H-dN) was reduced to 17.5 nM, the resultant chromosomes were comparable with those assembled with 35 nM holo(WT), in terms of both the morphology and the mSMC4 signal levels (Fig 1C and D). Taken together, we concluded that deletion of the CAP-H N-tail accelerates the loading of condensin I on chromosomes in M-HSS, and that the chromosomal levels of condensin I are tightly correlated with the chromosome morphology produced.

To gain further insights into the functional contribution of the CAP-H N-tail to condensin I-mediated chromosome assembly, we next focused on the conserved helix (CH), located in the middle of the N-tail, that is enriched with basic amino acids (Fig 1A). We constructed a holocomplex containing hCAP-H that lacks the CH, holo(H-dCH), and a holocomplex containing hCAP-H in which the basic amino acids within the CH are substituted with acidic residues, holo(H-crCH) (Figs 1A and EV1A). Here dCH and crCH stand for <u>d</u>eletion of the <u>CH</u> and <u>c</u>harge-<u>r</u>eversed <u>CH</u>, respectively. We found that the holo(H-dCH) and holo (H-crCH) exhibited a phenotype very similar to that of holo(H-dN) in terms of both increased condensin I loading and resultant chromosome morphology (Fig EV2A and B). Thus, the conserved helix and its positive charges play an important role in negatively regulating condensin I loading in M-HSS.

### Phosphorylation-deficient and phosphorylation-mimetic mutations of the CAP-H N-tail decelerate and accelerate condensin I loading, respectively

We next wished to understand how phosphorylation of the CAP-H N-tail might affect its function. Previous studies had shown that *Xenopus* CAP-H is phosphorylated in a mitosis-specific manner in *Xenopus* egg extracts (Hirano *et al*, 1997; Kimura *et al*, 1998), and that *Xenopus* and human CAP-H can be phosphorylated by cyclin B-Cdk1 *in vitro* (Kimura *et al*, 1998; Kimura *et al*, 2001; Shintomi *et al*, 2015). Whereas hCAP-H has three SP sites and one TP site in its N-tail (Fig EV3A), it is known that Cdk1 phosphorylation is not exclusively proline-directed (Brown *et al*, 2015; Suzuki *et al*, 2015; Krasinska & Fisher, 2022). Moreover, a phospho-proteomic analysis identified phosphorylation at both SP/TP and non-SP/TP sites in this region (Hornbeck *et al*, 2015). Thus, the targets of mitosis-specific phosphorylation in CAP-H have not yet been fully characterized.

To test whether phosphorylation of the hCAP-H N-tail plays an important role in condensin I loading, we prepared a holocomplexes harboring phosphorylation-deficient mutations in the CAP-H N-tail, holo(H-N19A), where all serines and threonines present in the N-tail were substituted with alanines (Figs 1A and EV1A). When the holo(WT) and holo(H-N19A) were incubated with M-HSS, two phosphoepitopes (pS17 and pS76) were detectable in the former, but not in the latter (Fig EV3A and EV3B; Materials and Methods). Neither of the epitopes was detected in both complexes that had been incubated with interphase HSS (I-HSS), indicating that the corresponding sites are phosphorylated in holo(WT) in a mitosis-specific manner. Holo(H-N19A) was then subjected to the add-back assay in M-HSS. We found that loading of holo(H-N19A) onto chromatin was greatly reduced compared to holo(WT) or holo(H-dN) (Fig 2A-C). The resultant chromosomes produced by holo(H-N19A) were poorly organized and abnormally thick, having hazy surfaces (Fig 2A-C).

**Figure 2.**
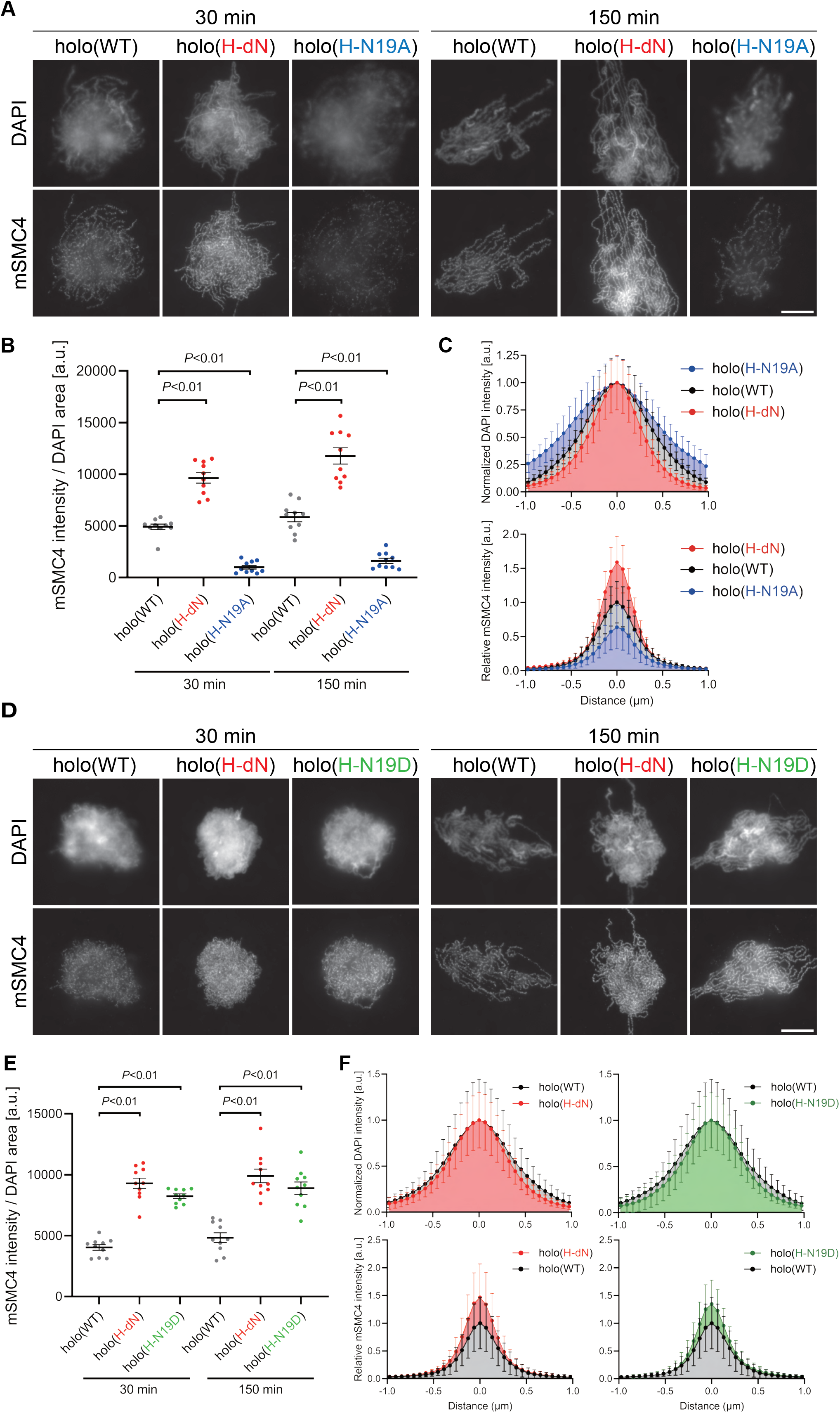
Phosphorylation-deficient and phosphorylation-mimetic mutations of the CAP-H N-tail decelerate and accelerate condensin I loading, respectively. (A) Mouse sperm nuclei were incubated with condensin-depleted M-HSS that had been supplemented with holo(WT), holo(H-dN), or holo(H-N19A) at a final concentration of 35 nM. After 30 min and 150 min, the reaction mixtures were fixed and processed for immunofluorescence labeling with an antibody against mSMC4. DNA was counterstained with DAPI. Scale bar, 10 μm. (B) Quantification of the intensity of mSMC4 per the DAPI area in the experiment shown in Fig 2A (n = 10 clusters of chromosomes). The error bars represent the mean ± SEM. The *P* values were assessed by Tukey’s multiple comparison test after obtaining a significant difference with one-way ANOVA at each time point. This experiment was reproduced more than three times for holo(WT), holo (H-dN), and holo(H-N19A). (C) Line profiles of mitotic chromosomes observed at 150 min in the experiment shown in Fig 2A. Signal intensities of DAPI (top) and mSMC4 (bottom) of the chromosomes assembled with holo(WT) (black), holo(H-dN) (red), or holo(H-N19A) (blue) were measured along with the lines drawn perpendicular to chromosome axes (n = 20). The mean and standard deviation were normalized individually to the DAPI intensities (arbitrary unit [a.u.]) at the center of chromosome axes (distance = 0 μm) within each set. Intensities of mSMC4 signals from holo(H-dN) and holo(H-N19A) were normalized relative to the value from holo(WT). (D) Add-back assay using holo(WT), holo(H-dN), and holo(H-N19D) at a final concentration of 35 nM was performed as described in Fig 2A. Scale bar, 10 μm. (E) Quantification of the intensity of mSMC4 per the DAPI area in the experiment shown in Fig 2D (n = 10 clusters of chromosomes). The error bars represent the mean ± SEM. The *P* values were assessed by Tukey’s multiple comparison test after obtaining a significant difference with one-way ANOVA at each time point. This experiment was reproduced more than three times for holo(WT) and holo (H-dN), and twice for holo(H-N19D). (F) Line profiles of mitotic chromosomes assembled observed at 150 min in the experiment shown in Fig 2D. The signal intensities of DAPI (top) and mSMC4 (bottom) of the chromosomes assembled with holo(WT) (black), holo(H-dN) (red), or holo(H-N19D) (green) were measured and plotted as described in Fig 2C (n = 20).

To further substantiate the observations described above, we then prepared and tested a holocomplex harboring phosphorylation-mimetic mutations, holo(H-N19D) (Figs 1A and EV1A), in which all serines and threonines in the N-tail were substituted with aspartic acids. Remarkably, we found that holo(H-N19D) behaved very similarly to holo(H-dN) in the add-back assay (Fig 2D-F). These results supported the idea that multisite phosphorylation in the CAP-H N-tail relieves its negative effects on condensin I loading and mitotic chromosome assembly.

### Deletion of the CAP-H N-tail enables condensin I to assemble mitotic chromosome-like structures even in interphase extracts

Having observed the accelerated loading of holo(H-dN) in M-HSS, we wondered how the mutant complex would behave in interphase extracts. To eliminate potential complications from the subcellular localization of condensin I (Ono *et al*, 2004; Shintomi & Hirano, 2011), we used membrane-free I-HSS in the following experiments in which no nuclear envelope was assembled around the chromatin introduced. Consistent with the previous study (Hirano *et al*, 1997), endogenous *Xenopus* condensins failed to associate with chromatin in I-HSS, leaving a spherical compact mass, although they bound to chromatin and assembled a cluster of rod-shaped chromosomes in M-HSS (Fig 3A). We then monitored the morphological changes of sperm nuclei in a condensin-depleted I-HSS. We found that the nuclei were transiently swollen at early time points (15-30 min) and then converted into compact chromatin masses at a late time point (150 min) (Fig 3B). The terminal morphology was very similar to that observed in the unperturbed I-HSS described above (Fig 3A). The same was true in the condensin-depleted extract that had been supplemented with 35 nM holo(WT). Signals of mSMC4 were hardly observed on the chromatin masses under this condition (Fig 3B). When the condensin-depleted extract was supplemented with 35 nM holo(H-dN), however, significant levels of mSMC4 signals were detectable on chromatin, especially at early time points (Fig 3D). It was also noticed that the compact masses observed at 150 min had somewhat irregular surfaces.

**Figure 3.**
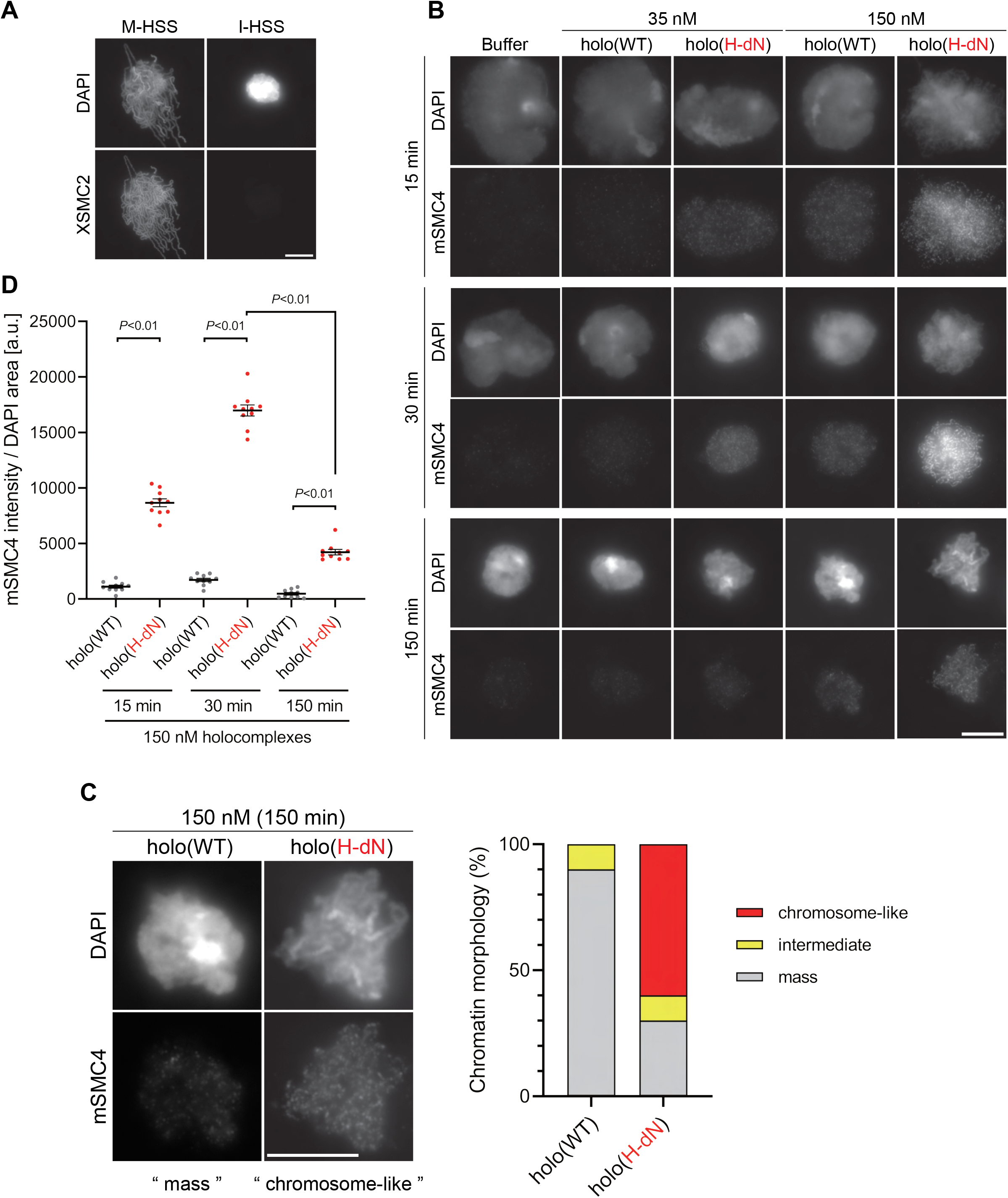
Deletion of the CAP-H N-tail enables condensin I to assemble mitotic chromosome-like structures even in interphase extracts. (A) Mouse sperm nuclei were incubated with M-HSS (left) and I-HSS (right) for 150 min, and the reaction mixtures were fixed and processed for immunofluorescence labeling with an antibody against XSMC2 (bottom). DNA was counterstained with DAPI (top). Scale bar, 10 μm. (B) Mouse sperm nuclei were incubated with condensin-depleted I-HSS that had been supplemented with a control buffer, holo(WT), or holo(H-dN) at a final concentration of 35 nM or 150 nM. After 15, 30 min, and 150 min, the reaction mixtures were fixed and processed for immunofluorescence labeling with an antibody against mSMC4. DNA was counterstained with DAPI. Scale bar, 10 μm. (C) Blow-up images of a chromatin “mass” assembled with 150 nM holo(WT) at 150 min and a cluster of mitotic “chromosome-like” structures assembled with 150 nM holo(H-dN) at 150 min in the experiment shown in Fig 3B. Frequencies of the chromosomal phenotypes observed under the two conditions, which include “intermediate” structures, are plotted on the right (n = 10 for each condition). Scale bar, 10 μm. (D) Quantification of mSMC4 intensities on interphase chromatin when added holo(WT) or holo(H-dN) at 150 nM. The graph shows the intensity of mSMC4 per the DAPI area (n = 10 masses of chromatin) shown in Fig 3B. The error bars represent the mean ± SEM. The *P* values were assessed by Tukey’s multiple comparison test after obtaining a significant difference with two-way ANOVA. This result was reproduced in three independent experiments, and another set of the reproduced result was also shown in Fig EV4.

To further extend the observations obtained at the standard concentration (35 nM) of condensin I added back into the extracts, we tested a higher concentration (150 nM) of holo(WT) and holo(H-dN) in the same assay. We found that 150 nM holo(WT) produced a temporal series of chromatin masses that was very similar to that observed with 35 nM holo(H-dN). In contrast, 150 nM holo(H-dN) produced morphological phenotypes strikingly different from the other conditions tested. At 15 and 30 min, high levels of mSMC4 signals were detectable on chromatin masses in which entangled thin chromatin fibers were observed (Fig 3B and D). At 150 min, although mSMC4 signals on chromatin decreased, a cluster of rod-shaped, mitotic chromosome-like structures with mSMC4-positive axes was clearly discernible in each of the chromatin masses (Fig 3B and D). A blow-up image of a representative example is shown in Fig 3C. It is important to note that not only the SP/TP sites in the CAP-H N-tail but also those in the CAP-D2 C-tail were barely phosphorylated in I-HSS (Fig EV3A and B).

Finally, holo(H-crCH) and holo(H-N19D) were subjected to the same add-back assay at the concentration of 150 nM (Fig EV4A and B). We found that both mutant complexes behaved similarly to holo(H-dN) although the loading efficiency of holo(H-N19D) was somewhat lower than that of holo(H-crCH). These results demonstrated that, when the CAP-H N-tail is compromised, condensin I gains the ability to assemble mitotic chromosome-like structures even in interphase extracts.

### Deletion of the CAP-H N-tail enhances topological loading onto circular DNA and increases the frequency of loop formation *in vitro*

We next compared the activities of holo(WT) and holo(H-dN) in two different functional assays without the aid of *Xenopus* egg extracts. In the first setup,we employed the so-called topological loading assay that had been originally developed to assess an ATP-dependent topological entrapment of nicked circular DNA by the cohesin ring (Murayama & Uhlmann, 2014). Briefly, holo(WT) and holo(H-dN) were incubated with nicked circular or linearized DNA in the absence of nucleotides or the presence of ATP or AMP-PNP, and then condensin-DNA complexes were recovered on beads by immunoprecipitation. After being washed with a high-salt buffer, the samples were split into two, and the DNA and proteins that remained on the beads were analyzed by agarose gel electrophoresis (Fig 4A) and immunoblotting (Fig 4B). Under this condition, virtually no linear DNA was recovered with either holo(WT) or holo(H-dN) regardless of the presence or absence of nucleotides (Fig 4A). In contrast, nicked circular DNA was recovered on the beads with holo(WT) in an ATP-dependent manner. Notably, a significantly higher amount (~2.5 fold) of nicked circular DNA was recovered with holo(H-dN) than with holo(WT) (Fig 4A and C).

**Figure 4.**
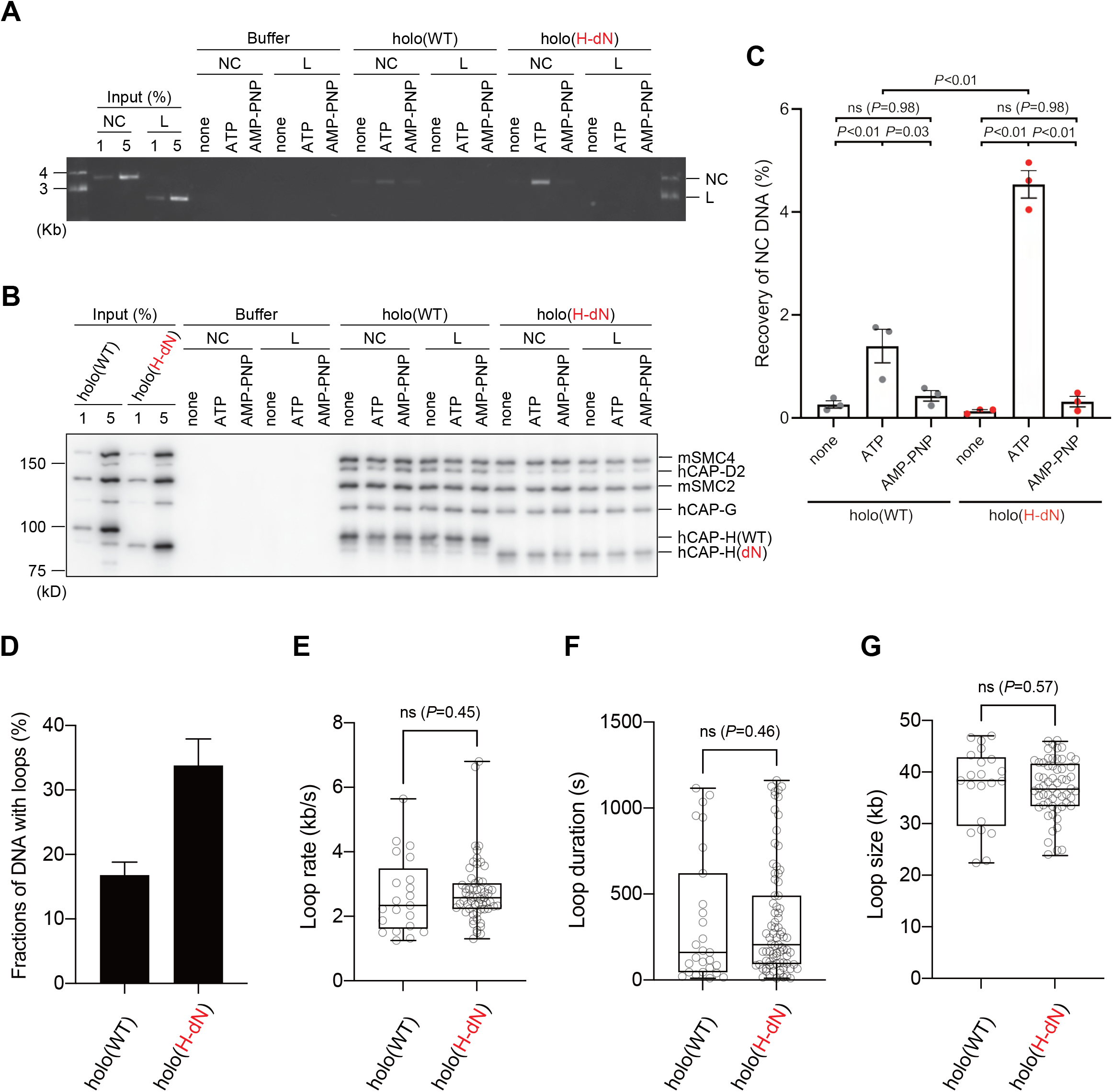
Deletion of the CAP-H N-tail enhances topological loading onto circular DNA and increases the frequency of loop formation *in vitro*. (A-C) Topological loading assay. (A) Loading reactions onto nicked circular DNA (NC) or linearized DNA (L) were set up with a control buffer, holo(WT) or holo(H-dN) in the absence of nucleotides (-) or the presence of ATP or AMP-PNP. DNAs recovered on beads after a high-salt wash were analyzed by agarose gel electrophoresis. (B) Confirmation of the efficiency of immunoprecipitation in the experiment shown in Fig 4A. Using the one-tenth volume of the recovered samples, immunoblotting analysis was performed with the antibodies indicated. (C) Quantification of the recovery of NC DNA in the experiments shown in Fig 4A. The error bars represent the mean ± SEM from three independent experiments. The *P* values were assessed by Tukey’s multiple comparison test after obtaining a significant difference with two-way ANOVA. (D-G) Loop extrusion assay. The error bars represent mean ± SEM. The *P* values shown in Fig 4E-G were assessed by a two-tailed Mann-Whitney *U* test. (D) Frequency of DNA loop formation by holo(WT) or holo(H-dN) (n = 3, ≥ 53 DNAs per condition). (E) Loop extrusion rate by holo(WT) or holo(H-dN) (from three independent experiments, n = 21 and 59 for holo(WT) and holo(H-dN), respectively). (F) Duration time to maintain DNA loops by holo(WT) or holo(H-dN) in the same experiment as in (E). (from three independent experiments, n = 27 and 81 for holo(WT) and holo(H-dN), respectively). (G) Loop size produced by holo(WT) or holo(H-dN) in the same experiment as in (E). (from three independent experiments, n = 21 and 59 for holo(WT) and holo(H-dN), respectively).

In the second setup, a loop extrusion assay using single DNA molecules was performed (Sakata *et al*, 2021; Kinoshita *et al*, 2022). In brief, holo(WT) and holo(H-dN) were labeled with Alexa Fluor 488 through a HaloTag (Fig EV1B) and subjected to the assay using U-shaped DNA. We found that holo(WT) and holo(H-dN) formed DNA loops on ~17% and ~33% of the U-shaped DNA, respectively, in the presence of ATP (Fig 4D). Despite the difference in the frequency of loop formation, no significant differences were observed between holo(WT) and holo(H-dN) in other parameters, such as the rate of loop extrusion, the loop duration time or the loop size formed (Fig 4E-G). Taken together, these results show that deletion of the CAP-H N-tail affects the initial interactions with DNA, but not the core activity responsible for the expansion of DNA loops.

### Concluding remarks and perspectives

In the current study, we have constructed and tested a panel of mutant complexes to provided evidence that the N-tail of the kleisin subunit CAP-H negatively regulates the loading of condensin I and the resultant assembly of mitotic chromosomes in *Xenopus* egg extracts (Fig 5A). Recent studies from our laboratory showed that deletion of the CAP-D2 C-tail, which also contains multiple SP/TP sites (Fig EV3A), has little impact on condensin I function as judged by the same and related add-back assays using *Xenopus* egg extracts (Kinoshita *et al*, 2022; Yoshida *et al*, 2022). Thus, the CAP-H N-tail represents the first example of negative regulatory elements that have been identified in vertebrate condensin I. Phosphorylation-deficient mutations (H-N19A) and phosphorylation-mimetic mutations (H-N19D) in the N-tail decelerate and accelerate condensin I loading, respectively, allowing us to propose the following working model (Fig 5B). The SMC-kleisin gate is closed when the CAP-H motif I binds to the SMC2 (Hassler *et al*, 2019) and prevents its untimely opening during interphase. The conserved helix located in the middle of the N-tail could play a critical contribution to this stabilization. Upon mitotic entry, multisite phosphorylation of the N-tail relieves the stabilization, allowing the opening of the DNA entry gate, hence, the loading of condensin I onto chromosomes. Thus, the kleisin N-tail of vertebrate condensin I could act as a “gatekeeper” of the SMC-kleisin gate. It is possible that, analogous to the Cdk1 inhibitor Sic1 (Örd *et al*, 2019), multisite phosphorylation of the CAP-H N-tail sets up an ultrasensitive switch-like response that can be activated at a certain threshold level of phosphorylation, thereby ensuring robust cell cycle regulation of condensin I loading.

**Figure 5.**
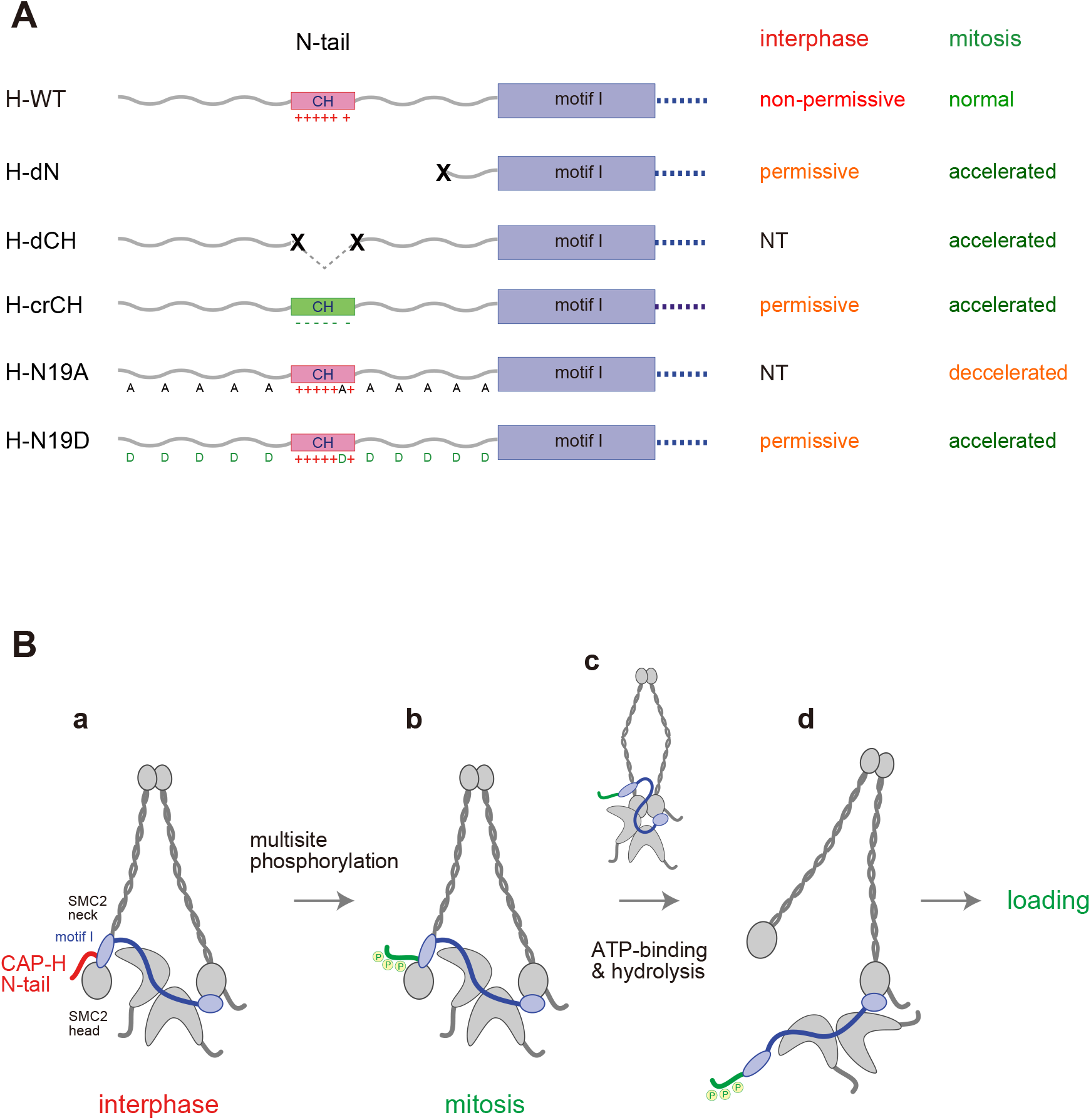
Regulation of condensin I by the N-tail of its kleisin subunit. (A) Summary of the mutant complexes tested in the current study. NT: not tested. (B) Model of cell cycle regulation of condensin I loading. The SMC-kleisin gate is closed when the CAP-H motif I binds to the SMC2 neck (a). In our model, this binding is stabilized by the CAP-H N-tail, thereby preventing its untimely opening during interphase. The conserved helix located in the middle of the N-tail contributes to this stabilization, possibly through direct interaction with the SMC2 head or neck. Upon mitotic entry, multisite phosphorylation of the N-tail relaxes this stabilization (b). ATP binding and hydrolysis by the SMC subunits trigger the opening of the DNA entry gate, thereby enabling condensin I to load onto chromosomes (c and d). Thus, the kleisin N-tail of vertebrate condensin I could act as a “gatekeeper” of the SMC-kleisin gate.

Our results show that, when the CAP-H N-tail function is compromised, the mutant condensin I complexes gain the ability to assemble mitotic chromosome-like structures even in interphase extracts. This raises the intriguing possibility that vertebrate condensin I can escape the tight requirement for Cdk1 phosphorylation under certain conditions. Finally, it should be mentioned that, although our recombinant condensin I complexes work excellently in the add-back assays using *Xenopus* egg extracts (Kinoshita *et al*, 2015; Kinoshita *et al*, 2022)(the current study), we have so far been unsuccessful in using these recombinant complexes to recapitulate positive DNA supercoiling or chromatid reconstitution, both of which require proper Cdk1 phosphorylation *in vitro* (Kimura *et al*, 1998; Shintomi *et al*, 2015). Faithfully reconstituting the key activities of condensin I using the recombinant subunits and dissecting multiple phosphorylation events required for its activation are important issues to be addressed in the future.

## Materials and Methods

### Preparation of recombinant condensin I complexes

DNA constructs for expressing condensin I subunits in insect cells were described previously (Kinoshita *et al*, 2022). Among them, a special note on the hCAP-H sequence is placed here because multiple cDNA sequences encoding different isoforms of hCAP-H with different lengths have been deposited in the database. Among them, the longest annotated polypeptide of hCAP-H is 741 amino-acid long (NP_056156.2). For historical reasons, our laboratory has been using a sequence of 730 amino-acid long as our reference sequence, whose translation starts at the second methionine of NP_056156.2 (Onn *et al*, 2007; Kinoshita *et al*, 2015; Kinoshita *et al*, 2022). The sequence corresponding to the first 11 amino acids of NP_056156.2 is completely missing in the ortholog of *Xenopus laevis*, and is not conserved in the ortholog of *Mus musculus* (in the latter case, the corresponding sequence is 8 amino-acid long rather than 11 amino-acid long). It is therefore very unlikely that the polypeptide of 730-amino-acid long used in the current study behaves differently from that of 741-amino-acid long. For this reason, the sequence of 730 amino-acid long is used as a “full-length” hCAP-H in the current study. The recombinant expression vector containing mSMC2 and mSMC4 cDNAs was described previously (Kinoshita *et al*, 2015). The cDNA sequences for hCAP-D2, -G and -H codon-optimized for the cabbage looper *Trichoplusia ni* were synthesized (GeneArt Gene Synthesis Service) and cloned into the pFastBac1 vector (Thermo Fisher Scientific). Deletion of the predicted helix motif in the hCAP-H N-tail was performed using PrimeSTAR Mutagenesis Basal Kit (TaKaRa). Primers used in the deletion were as follows: forward 5’-GAACGACTTCTCTACCGACTCTCCC-3’, reverse 5’-GTAGAGAAGTCGTTCTGAGGGAAGTC-3’. Full-length and N-terminally deleted versions of HaloTag-hCAP-H vectors were constructed by fusing the 3’ ends of the corresponding cDNAs with a HaloTag fragment using In-fusion HD Cloning Kit (TaKaRa). The following primers were used the construction of these HaloTag-hCAP-H vectors: hCAP-H (WT and dN) forward 5’-GAAGCGCGCGGAATTCGCCA-3’, hCAP-H (WT) reverse 5’-GAAGTACAGGTCCTCAGTGGTAGGTTCCAGGTCGCCCTGCCTAACTAA-3’, hCAP-H (dN) reverse 5’-GAAGTACAGGTCCTCAGTGGTAGGTTCCAGGTCGCCCTGCCTGACTAA-3’, HaloTag forward 5’-GAGGACCTGTACTTCCAGTCTGACAACGACATGGCCGAAATCGGAACT-3’ HaloTag reverse 5’-AGCGGCCGCGACTAGTTTATCCGCTGATTTCCAGGGTA-3’. DH10Bac (Thermo Fisher Scientific) was transformed with the plasmid vectors to produce bacmid DNAs, with which Sf9 cells were transfected to generate the corresponding baculoviruses. To produce recombinant condensin I complexes, High Five cells (Thermo Fisher Scientific) were infected with the baculoviruses carrying the condensin I subunits, and the resultant complexes were purified from cell lysates by two-step column chromatography as described previously (Kinoshita *et al*, 2022).

### Antibodies

Primary antibodies used in the present study were as follows: anti-XSMC4 (in-house identifier AfR8L, affinity-purified rabbit antibody), anti-XSMC2 (AfR9-6 for immunodepletion and immunoblotting, biotin-labeled AfR9-6 for immunofluorescence, affinity-purified rabbit antibody), anti-XCAP-D2 (AfR16L, affinity-purified rabbit antibody), anti-XCAP-G (AfR11-3L, affinity-purified rabbit antibody), anti-XCAP-H (AfR18, affinity-purified rabbit antibody; Hirano & Mitchison, 1994; Hirano *et al*, 1997), anti-XCAP-D3 (AfR192-2L, affinity-purified rabbit antibody), anti-XCAP-H2 (AfR201-4L, affinity-purified rabbit antibody; Ono *et al*, 2003), anti-mSMC4 (AfR326-3L, affinity-purified rabbit antibody), anti-mSMC2 (AfR329-4L, affinity-purified rabbit antibody; Lee *et al*, 2011), anti-hCAP-D2 (AfR51-3, affinity-purified rabbit antibody), anti-hCAP-G (AfR55-4, affinity-purified rabbit antibody), anti-hCAP-H (AfR57-4, affinity-purified rabbit antibody; Kimura *et al*, 2001), anti-XTopo IIα (αC1-6, rabbit antiserum; Hirano & Mitchison, 1993). Two custom phospho-specific antibodies against hCAP-H (AfR464-3P and AfR470-3P, affinity-purified rabbit antibodies) were raised against the phosphopeptides hHP1 ([C]PHSASpSPSERV) and hHP2 ([C]PRLLApSPSSRS), respectively (SIGMA Genosys), where cysteines (C) were added for chemical conjugation. The phosphoserines in hHP1 and hHP2 corresponded to pS17 and pS76 in the 730 amino-acid long isoform, respectively, and to pS28 and pS87 in the 741 amino-acid long isoform, respectively (see above for the explanation of the isotypes with different lengths). Three phospho-specific antibodies against hCAP-D2 (AfR173-4P, AfR175-4P and AfR177-4P, affinity-purified rabbit antibodies) were raised against the phosphopeptides hDP1 ([C]DNDFVpTPEPRR), hDP2 ([C]MTEDEpTPKKTT), and hDP3 ([C]TPKKTpTPILRA), respectively, and purified according to the procedure described previously (Kimura *et al*, 1998). The phosphothreonines in hDP1, hDP2 and hDP3 corresponded to pT1339, pT1384, and pT1389, respectively (Fig EV3A). Secondary antibodies used in the present study were as follows: Alexa Fluor 568-conjugated anti-rabbit IgG (A11036 [RRID: AB_10563566], Thermo Fisher Scientific), Alexa Fluor 488-conjugated anti-mouse IgG (A11001 [RRID: AB_2534069], Thermo Fisher Scientific), Alexa Fluor 488-conjugated streptavidin (S11223, Thermo Fisher Scientific), and horseradish peroxidase-conjugated anti-rabbit IgG (PI-1000 [RRID: AB_2336198], Vector Laboratories).

### Animals

Female *Xenopus laevis* frogs (RRID: NXR 0.031, Hamamatsu Seibutsu-Kyozai) were used to lay eggs to harvest *Xenopus* egg extract (Hirano *et al*, 1997). Frogs were used in compliance with the institutional regulations of the RIKEN Wako Campus. Mice (BALB/c × C57BL/6J)F1) for sperm nuclei (Shintomi *et al*, 2017) were used in compliance with protocols approved by the Animal Care and Use Committee of the University of Tokyo (for M. Ohsugi who provided mouse sperm).

### Preparation of *Xenopus* egg extracts

The high-speed supernatant of metaphase-arrested *Xenopus* egg extracts (M-HSS) was prepared as described previously (Shintomi & Hirano, 2018). For the preparation of interphase HSS, a low-speed supernatant (LSS) of metaphase-arrested extracts was supplemented with CaCl_2_ and cycloheximide at final concentrations of 0.4 mM and 100 μg/ml, respectively (Losada *et al*, 2002). After incubation at 22°C for 30 min, the interphase LSS was further fractionated by centrifugation at 50,000 rpm at 4°C for 90 min to yield interphase HSS (I-HSS).

### Immunodepletion, add-back assay and immunofluorescence

Immunodepletion of endogenous *Xenopus* condensin I and II from the extracts was performed using Dynabeads Protein A (Thermo Fisher Scientific) as described previously (Δcond) (Kinoshita *et al*, 2022). Add-back assays using mouse sperm nuclei were performed as described previously (Kinoshita *et al*, 2022). After incubation for indicated time points, the reaction mixtures were fixed with 10 volumes of KMH (20 mM K-HEPES [pH 7.7], 100 mM KCl, and 2.5 mM MgCl_2_) containing 4% formaldehyde and 0.1% Triton X-100 at 22°C for 15 min. The fixed chromatin was sedimented onto a coverslip through 5-ml cushion of XBE2 (10 mM K-HEPES [pH 7.7], 100 mM KCl, 2 mM MgCl_2_, 0.1 mM CaCl_2_, 5 mM EGTA, and 50 mM sucrose) containing 30% glycerol by centrifugation at 5,000 rpm for 10 min. For immunofluorescence, the coverslips were blocked with TBS-Tx containing 1% BSA for 30 min and then incubated with primary antibodies for 60 min. After being washed three times with TBS-Tx, the coverslips were incubated with secondary antibodies for 60 min. After DNA was stained with DAPI, they were mounted with VectaShield mounting medium (H-1000, Vector Laboratories) onto slides and then observed under an Olympus BX63 fluorescent microscope equipped with a UPlanSApo 100×/1.40 NA oil immersion lens and an ORCA-Flash 4.0 digital complementary metal oxide semiconductor camera C11440 (Hamamatsu Photonics). The fluorescent images were acquired using CellSens Dimension software (Olympus).

### Quantitative analyses of immunofluorescent images and statistics

Immunofluorescent images acquired by microscopic observation were quantified using ImageJ software (https://imagej.nih.gov/ij). Line profiles of mitotic chromosomes were scanned according to the procedure described previously (Kinoshita *et al*, 2022). For quantification of recombinant condensin I complexes on chromatin, the DAPI- and mSMC4-positive signals from the images were segmented using the threshold function and the integrated densities and the areas were measured. The mSMC4 intensities were divided by the DAPI-positive areas to assess the accumulation of condensin complexes on chromosome clusters. The data were handled with Excel software (Microsoft), and the graphs were drawn using GraphPad Prism software (version 8, GraphPad Software). Tukey’s multiple comparison test after obtaining a significant difference with one-way ANOVA (Fig 1B and E) or two-way ANOVA (Figs 1C, 3D, and EV4B).

### Immunoblotting

Denatured protein samples were subjected to SDS-PAGE and transferred onto a nitrocellulose membrane (Cytiva). The membrane was blocked with 5% skim milk in TBS-Tw (20 mM Tris-HCl [pH7.5], 150 mM NaCl, and 0.05% Tween 20) for 30 min and incubated with primary antibodies diluted with 1% BSA in TBS-Tw for 1 h. After being washed three times with TBS-Tw, the membrane was incubated with a peroxidase-conjugated secondary antibody diluted with TBS-Tw for 1 h. After being washed three times with TBS-Tw, the membrane was treated with a chemiluminescence substrate for peroxidase (WBKLS500, Merck). The chemiluminescent image was acquired using an image analyzer (Amersham Imager 680, Cytiva).

### Detection of phosphoepitopes in *Xenopus* egg extracts

For immunoprecipitation, rProtein A Sepharose (rPAS, Cytiva) was equilibrated with TBS. 5 μl of rPAS were coupled with 5 μg anti-hCAP-G antibody at 4°C for 2 h with rotation and then washed three times with KMH. Holo(WT) or holo(H-N19A) was added to Δcond M-HSS or I-HSS at a final concentration of 35 nM, along with 1/50 vol of 50× energy mix (50 mM Mg-ATP, 500 mM phosphocreatine, and 2.5 mg/ml creatine kinase [pH 7.5]) and incubated at 22°C for 150 min. The final reaction volume was 15 μl. The reaction mixtures were immunoprecipitated with the anti-hCAP-G antibody-coupled rPAS on ice for 30 min. The beads were washed with 150 μl KMH containing 20 mM β-glycerophosphate and 0.1% Triton X-100, and then centrifuged (7,000 rpm) at 4°C for 1 min. The recovered beads were further washed three times with 200 μl KMH containing 20 mM ß-glycerophosphate and 0.1% Triton X-100, followed by two repeated washes with 200 μl KMH containing 20 mM β-glycerophosphate. The immunoprecipitants were analyzed by immunoblotting.

### Topological loading assay

A topological loading assay was performed according to the procedure described by Murayama and Uhlmann (Murayama & Uhlmann, 2014) with minor modifications. Nicked circular DNA and linearized DNA were used as DNA substrates in this assay. To prepare the nicked circular DNA, negatively supercoiled plasmid DNA (pUC19) purified from *Escherichia coli* cells using Nucleobond PC100 (MACHERREY-NAGEL GmbH & Co. KG) was treated with the nicking enzyme Nt.BspQI (R0644S, New England Biolabs). The linearized DNA was prepared by digesting the negatively supercoiled DNA with *Eco* RI (1040A, TaKaRa Bio). The treated DNAs were extracted with phenol/chloroform/isoamyl alcohol (25:24:1) and then purified by ethanol precipitation. For condensin I loading assay, 2 pmol of holo(WT) or holo(H-dN) and 90 ng of nicked or linearized DNA were mixed in reaction buffer (35 mM Tris-HCl [pH 7.5], 1 mM TCEP, 50 mM NaCl, 7 mM MgCl_2_, 0.003% Tween 20, and 15% glycerol) and then supplemented with ATP or AMP-PNP (NU-407, Jena Bioscience) at a final concentration of 2 mM. The final reaction volume was 15 μl. The mixture was incubated at 32°C for 1 h. Note that the final concentration of NaCl in the loading reaction was 77 mM because of carry-over from the storage buffer of purified condensin I. The loading reaction was terminated by adding 85 μl of IP buffer (35 mM Tris-HCl [pH7.5], 0.5 mM TCEP, 500 mM NaCl, 10 mM EDTA, 5% glycerol, and 0.35% Triton X-100). Condensin I-DNA complexes were then immunoprecipitated with 10 μl of Dynabeads protein A (10002D, Thermo Fisher Scientific) that had been pre-coupled with 2.5 μg of affinity-purified anti-hCAP-G. The beads were washed three times with 750 μl of Wash Buffer-I (35 mM Tris-HCl [pH7.5], 0.5 mM TCEP, 750 mM NaCl, 10 mM EDTA, and 0.35% Triton X-100), and then once with 500 μl of Wash Buffer-II (35 mM Tris-HCl [pH7.5], 0.5 mM TCEP, 100 mM NaCl, and 0.1% Triton X-100). To check the efficiency of immunoprecipitation, protein samples were retrieved from a one-tenth volume of the beads and analyzed by immunoblotting. The remaining beads were treated with 15 μl of deproteinization solution (10 mM Tris-HCl [pH7.5], 1 mM EDTA, 50 mM NaCl, 0.75% SDS, and 1 mg/ml proteinase K) and incubated at 50°C for 20 min. The recovered DNAs were electrophoresed on a 1% agarose gel in TAE, and stained with GelRed (Biotium). The fluorescent images were acquired using an image analyzer (Amersham Imager 680, Cytiva). The signal intensities of nicked circular DNAs recovered by holo(WT) and holo(H-dN) were quantified using ImageJ software. Three independent experiments were performed and assessed by Tukey’s multiple comparison test after obtaining a significant difference with two-way ANOVA.

### Loop extrusion assay

The HaloTag-holocomplexes were labeled with HaloTag Alexa Fluor 488 Ligand (Promega) at room temperature for 30 min. The labeled complexes were applied onto a PD-10 column (Cytiva) to remove unbound fluorescent ligands. A loop extrusion assay was performed as described previously (Sakata *et al*, 2021; Kinoshita *et al*, 2022). In brief, coverslips were coated with 1 mg/ml streptavidin in ELB++ (10 mM HEPES-KOH [pH 7.7], 50 mM KCl, 2.5 mM MgCl_2_, and 1 mg/ml BSA) for 30 min. After assembling microfluidic flow cells that can switch the flow direction, they were incubated with ELB++ for 30 min, and then washed with T20 buffer (40 mM Tris-HCl [pH 7.5], 20 mM NaCl, and 0.2 mM EDTA). Biotin-labeled λ DNA in T20 buffer was attached to the coverslips for 5 min, and then unbound DNA was washed off with imaging buffer (50 mM Tris-HCl [pH 7.5], 50 mM NaCl, 5 mM MgCl_2_, 3 mM ATP, and 1 mM DTT). The flow direction was switched, and the flow cells were washed with the same buffer for another 10 min. Then, one nM condensin in the imaging buffer was introduced into the flow cells at 20 μl/min for 1.5 min, and washed off with the same buffer at 20 μl/min. For visualization of Sytox Orange (22 nM)-staining DNA and Alexa 488-labeled condensin, 488 nm and 561 nm lasers were used, respectively. Images were taken every 5 sec for 20 min after introducing condensin. The images were analyzed using the ImageJ software. The data in Fig 4E-G were statistically analyzed by a two-tailed Mann-Whitney *U* test.

## Materials availability

All unique/stable reagents generated in this study are available from the corresponding author.

## Acknowledgements

We thank F Inoue, H Watanabe, and M Ohsugi for their help with mouse sperm nuclei preparation, and members of the Hirano lab for critically reading the manuscript. This work was supported by Grant-in-Aid for Scientific Research/KAKENHI (#17K15070 [to ST], #19H05755 and #22H02551 [to KS], #19K06499 [to KK], #20H05937 [to TN], and #18H05276 and #20H0593 [to TH]) and by JST PRESTO (JPMJPRK4 [to TN]).

## Author contributions

ST and TH conceptualized the projects and designed the experiments. ST performed all experiments and corresponding image analyses except for the loop extrusion assay. KS prepared mouse sperm nuclei and provided technical instruction and advice to ST. KK instructed ST in the expression, purification, and fluorescence labeling of recombinant condensin I complexes. YT performed the loop extrusion assay and data analysis. TN supervised the loop extrusion assay. TH designed and purified the hCAP-D2 phospho-specific antibodies. ST and TH wrote the manuscript with input from all of the other authors.

## Supporting Information

Expanded View Figures PDF

## Expanded view figure legends

**Figure EV1.**
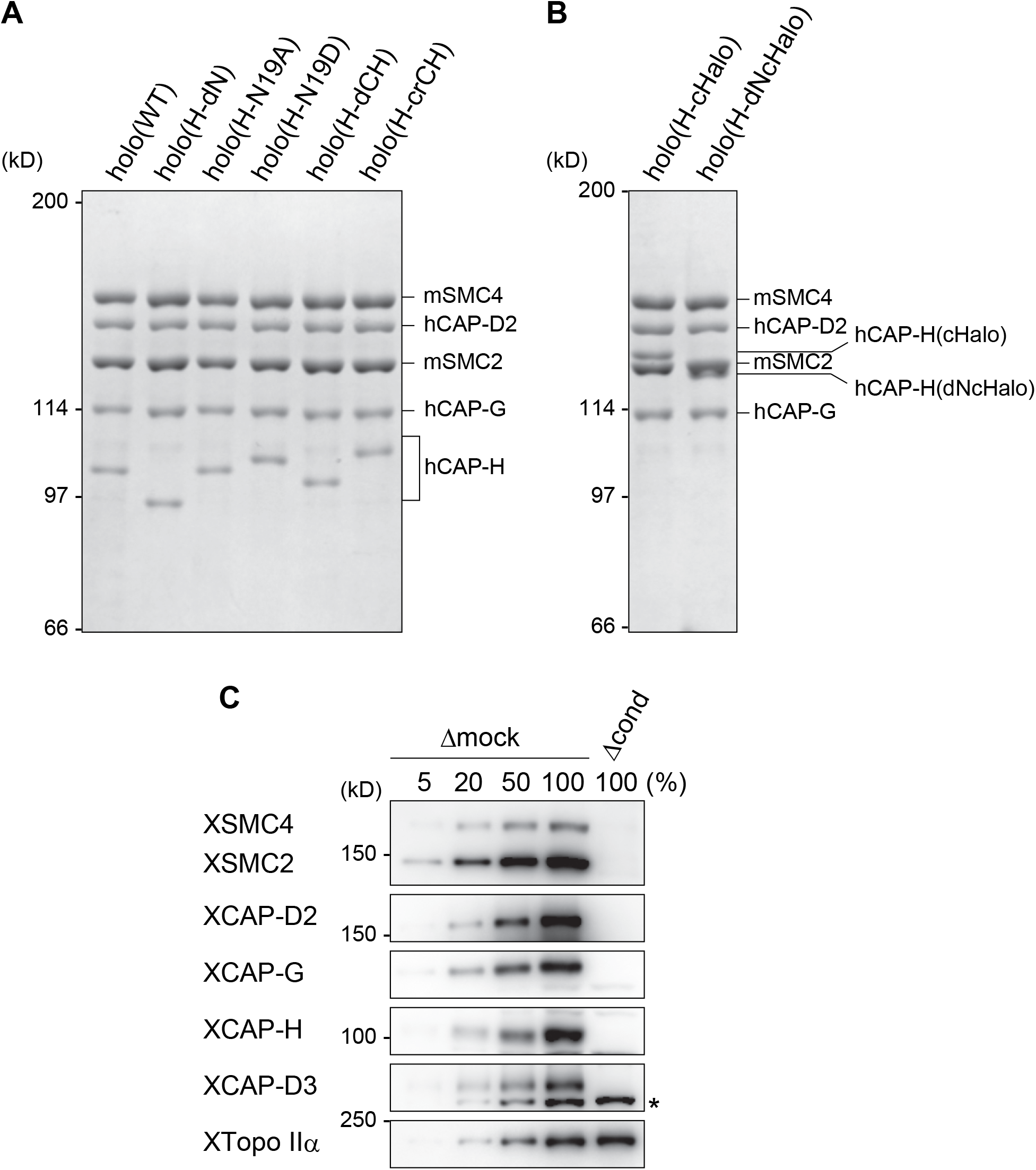
Recombinant condensin I complexes and immunodepletion. (A) The wild-type and mutant condensin I complexes used in the current study. The recombinant complexes were purified from insect cells and subjected to SDS-PAGE. The gel was stained with Coomassie brilliant blue (CBB). (B) HaloTag-condensin I complexes used in the loop extrusion assay. The recombinant complexes were purified from insect cells, fluorescence-labeled and subjected to SDS-PAGE. The gel was stained with CBB. (C) Immunodepletion of endogenous condensins from *Xenopus* egg M-HSS. Endogenous condensin subunits were depleted from the M-HSS using Dynabeads Protein A coupled with control IgG (Δmock) or a mixture of antibodies against the subunits of condensins I and II (Δcond). To estimate the efficiency of depletion, the Δcond extract (100%) was compared with different amounts of the Δmock extract (5, 20, 50 and 100%) by immunoblotting using the antibodies indicated. Endogenous topo IIα (XTopo IIα) was used as a loading control. The asterisk indicates a non-specific band.

**Figure EV2.**
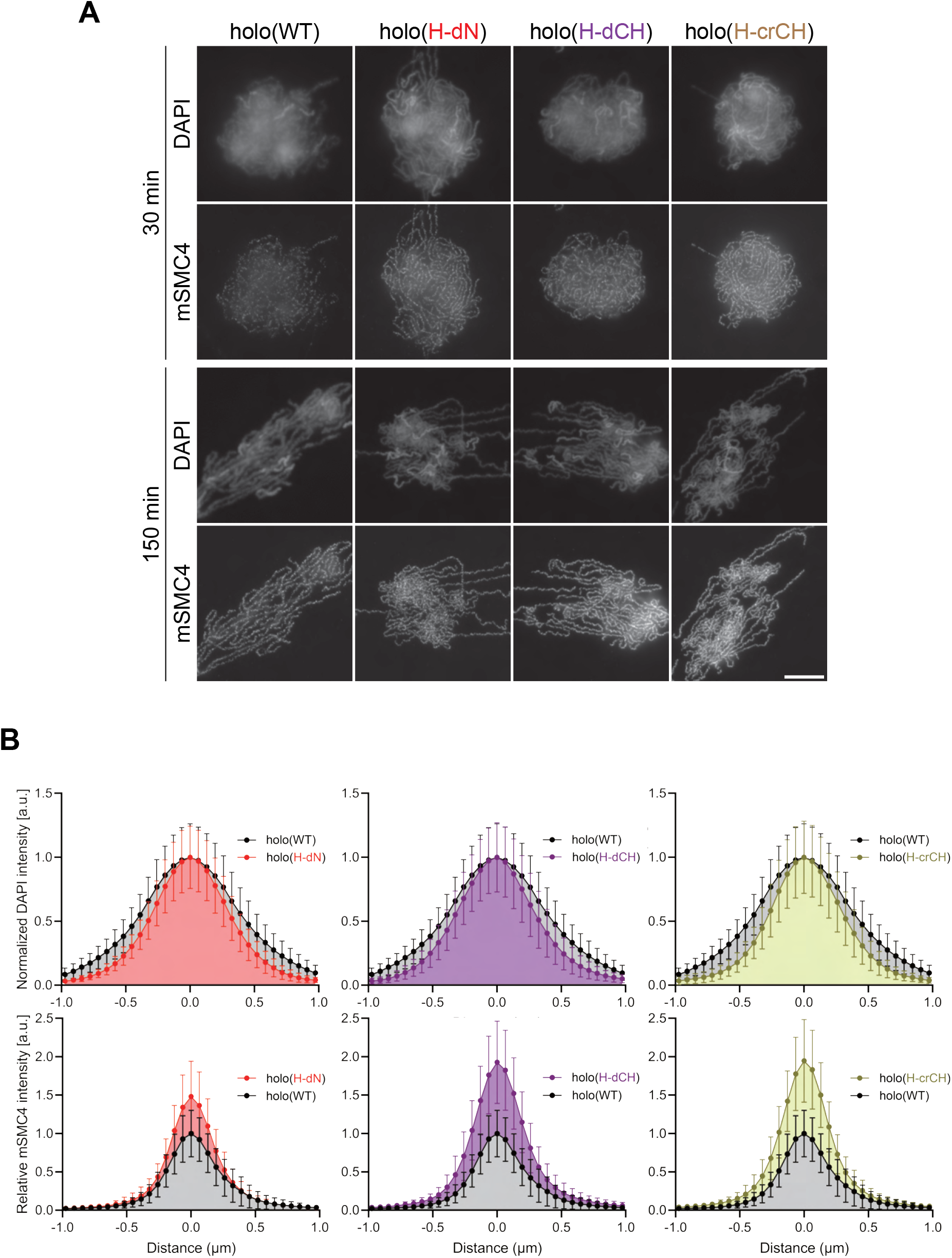
Deletion of and mutations in the conserved helix accelerates condensin I loading and mitotic chromosome assembly. (A) Mouse sperm nuclei were incubated with condensin-depleted M-HSS that had been supplemented with holo(WT), holo(H-dN) holo(H-dCH) or holo(H-crCH) at a final concentration of 35 nM. After 30 and 150 min, the reaction mixtures were fixed and processed for immunofluorescence labeling with an antibody against mSMC4. DNA was counterstained with DAPI. Scale bar, 10 μm. (B) Line profiles of mitotic chromosomes observed at 150 min in the experiment shown in Fig EV2A. Signal intensities of DAPI (top) and mSMC4 (bottom) from the chromosomes assembled with holo (WT) (black), holo(H-dN) (red), holo(H-dCH) (purple) or holo(H-crCH) (yellow) were measured along with the lines drawn perpendicular to chromosome axes (n = 20). The error bars represent the mean ± SD. The mean and standard deviation were normalized individually to the DAPI intensities (arbitrary unit [a.u.]) at the center of chromosome axes (distance = 0 μm) within each set. Intensities of mSMC4 signals from holo(H-dN), holo(H-dCH) and holo(H-crCH) were normalized relative to the value from holo(WT).

**Figure EV3.**
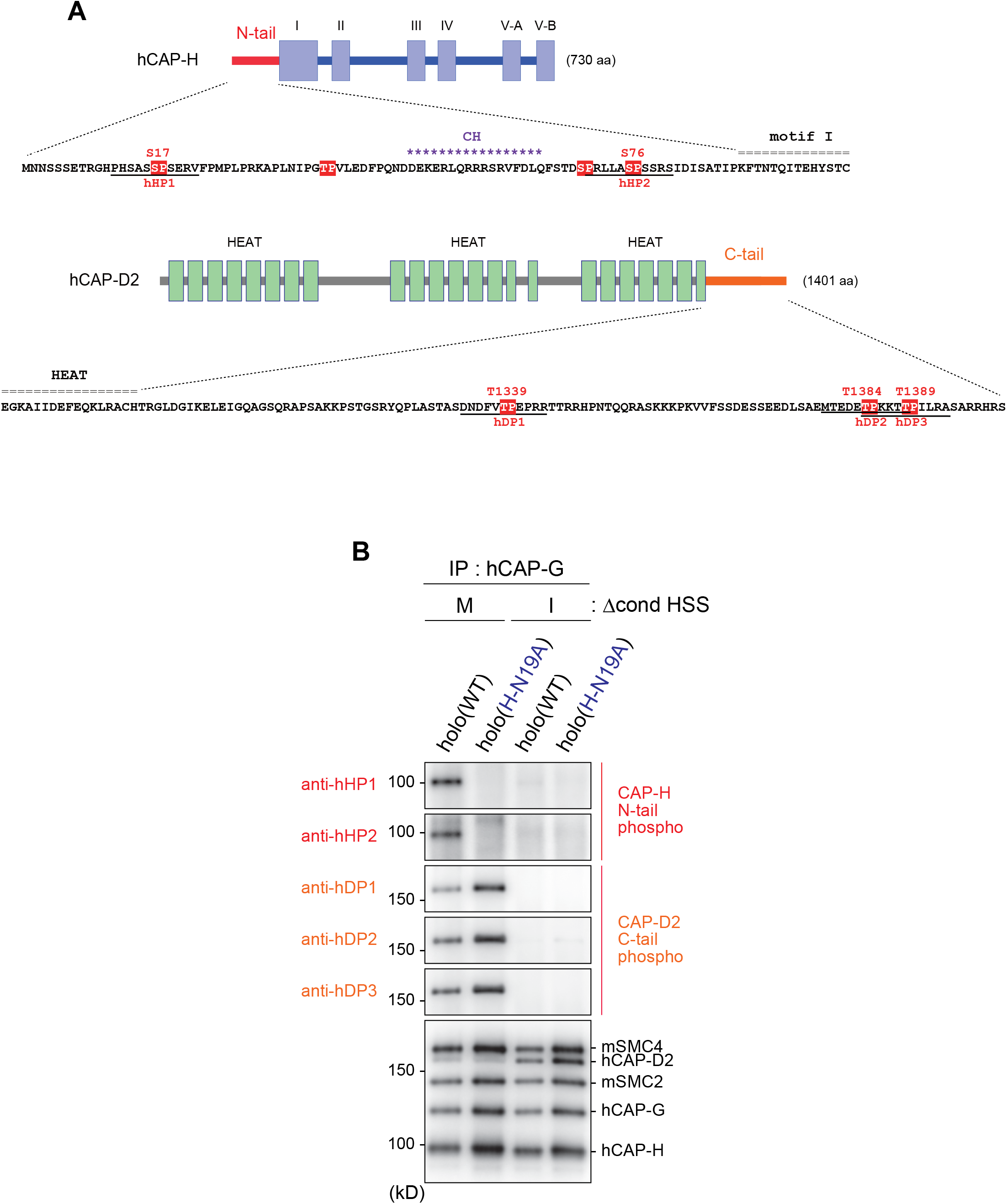
Mitosis-specific phosphorylation of condensin subunits. (A) Schematic diagrams of hCAP-H and hCAP-D2. The hCAP-H N-tail has four SP/TP sites whereas the hCAP-D2 C-tail has three TP sites. Peptide sequences used to prepare phospho-specific antibodies are underlined (hHP1 and hHP4 for hCAP-H; hDP1, hDP2 and hDP3 for hCAP-D2). (B) Detection of phophoepitopes in *Xenopus* egg extracts. Holo(WT) or holo(H-N19A) was added to Δcond M-HSS or I-HSS at a final concentration of 35 nM and incubated at 22°C for 150 min. The condensin complexes were immunoprecipitated and analyzed by immunoblotting using the antibodies indicated.

**Figure EV4.**
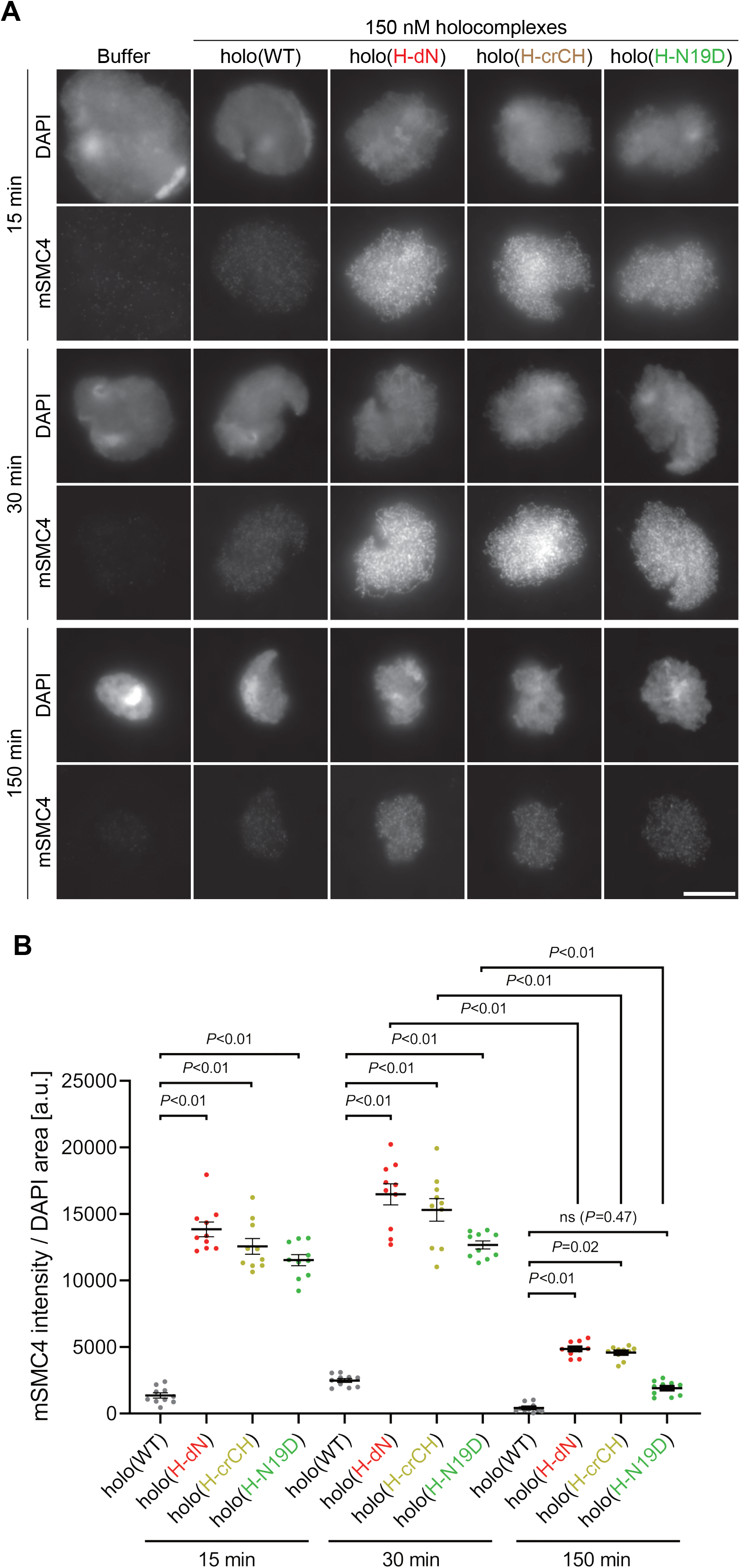
Deletion of and mutations in the conserved helix enables condensin I to assemble mitotic chromosome-like structures even in interphase extracts. (A) Mouse sperm nuclei were incubated with condensin-depleted I-HSS that had been supplemented with holo(WT), holo(H-dN), holo(H-crCH) or holo(H-N19D) at a final concentration of 150 nM. After 15, 30 and 150 min, the reaction mixtures were fixed and processed for immunofluorescence labeling with an antibody against mSMC4. DNA was counterstained with DAPI. Scale bar, 10 μm. (B) Quantification of the intensity of mSMC4 per DAPI area in the experiment shown in Fig EV4A (n = 10 clusters of chromosomes). The error bars represent the mean ± SEM. The *P* values were assessed by Tukey’s multiple comparison test after obtaining a significant difference with two-way ANOVA.

## References

Batty P, Gerlich DW (2019) Mitotic Chromosome Mechanics: How Cells Segregate Their Genome. Trends Cell Biol 29: 717–726

Bazile F, St-Pierre J, D’Amours D (2010) Three-step model for condensin activation during mitotic chromosome condensation. Cell Cycle 9: 3243–3255

Brown NR, Korolchuk S, Martin MP, Stanley WA, Moukhametzianov R, Noble MEM, Endicott JA (2015) CDK1 structures reveal conserved and unique features of the essential cell cycle CDK. Nat Commun 6: 6769

Gibcus JH, Samejima K, Goloborodko A, Samejima I, Naumova N, Nuebler J, Kanemaki MT, Xie L, Paulson JR, Earnshaw WC et al (2018) A pathway for mitotic chromosome formation. Science 359: eaao6135

Green LC, Kalitsis P, Chang TM, Cipetic M, Kim JH, Marshall O, Turnbull L, Whitchurch CB, Vagnarelli P, Samejima K et al (2012) Contrasting roles of condensin I and condensin II in mitotic chromosome formation. J Cell Sci 125: 15911604

Hara K, Kinoshita K, Migita T, Murakami K, Shimizu K, Takeuchi K, Hirano T, Hashimoto H (2019) Structural basis of HEAT-kleisin interactions in the human condensin I subcomplex. EMBO Rep 20: e47183

Hassler M, Shaltiel IA, Kschonsak M, Simon B, Merkel F, Thärichen L, Bailey HJ, Macošek J, Bravo S, Metz J et al (2019) Structural Basis of an Asymmetric Condensin ATPase Cycle. Mol Cell 74: 1175–1188.e9

Hirano T (2016) Condensin-Based Chromosome Organization from Bacteria to Vertebrates. Cell 164: 847–857

Hirano T, Kobayashi R, Hirano M (1997) Condensins, chromosome condensation protein complexes containing XCAP-C, XCAP-E and a Xenopus homolog of the Drosophila Barren protein. Cell 89: 511–521

Hirano T, Mitchison TJ (1993) Topoisomerase II does not play a scaffolding role in the organization of mitotic chromosomes assembled in Xenopus egg extracts. J Cell Biol 120:601–612

Hirano T, Mitchison TJ (1994) A heterodimeric coiled-coil protein required for mitotic chromosome condensation in vitro. Cell 79: 449–458

Hirota T, Gerlich D, Koch B, Ellenberg J, Peters JM (2004) Distinct functions of condensin I and II in mitotic chromosome assembly. J Cell Sci 117: 6435–6445

Holt LJ, Tuch BB, Villen J, Johnson AD, Gygi SP, Morgan DO (2009) Global analysis of Cdk1 substrate phosphorylation sites provides insights into evolution. Science 325:1682–1686

Hornbeck PV, Zhang B, Murray B, Kornhauser JM, Latham V, Skrzypek E (2015) PhosphoSitePlus, 2014: mutations, PTMs and recalibrations. Nucleic Acids Res 43: D512–520

Kimura K, Cuvier O, Hirano T (2001) Chromosome condensation by a human condensin complex in Xenopus egg extracts. J Biol Chem 276: 5417–5420

Kimura K, Hirano M, Kobayashi R, Hirano T (1998) Phosphorylation and activation of 13S condensin by Cdc2 in vitro. Science 282: 487–490

Kinoshita K, Kobayashi TJ, Hirano T (2015) Balancing acts of two HEAT subunits of condensin I support dynamic assembly of chromosome axes. Dev Cell 33: 94–106

Kinoshita K, Tsubota Y, Tane S, Aizawa Y, Sakata R, Takeuchi K, Shintomi K, Nishiyama T, Hirano T (2022) A loop extrusion-independent mechanism contributes to condensin I-mediated chromosome shaping. J Cell Biol 221: e202109016

Krasinska L, Fisher D (2022) A Mechanistic Model for Cell Cycle Control in Which CDKs Act as Switches of Disordered Protein Phase Separation. Cells 11: 2189

Kschonsak M, Merkel F, Bisht S, Metz J, Rybin V, Hassler M, Haering CH (2017) Structural Basis for a Safety-Belt Mechanism That Anchors Condensin to Chromosomes. Cell 171: 588–600.e24

Lee J, Ogushi S, Saitou M, Hirano T (2011) Condensins I and II are essential for construction of bivalent chromosomes in mouse oocytes. Mol Biol Cell 22: 34653477

Lipp JJ, Hirota T, Poser I, Peters JM (2007) Aurora B controls the association of condensin I but not condensin II with mitotic chromosomes. J Cell Sci 120: 12451255

Losada A, Hirano M, Hirano T (2002) Cohesin release is required for sister chromatid resolution, but not for condensin-mediated compaction, at the onset of mitosis. Genes Dev 16: 3004–3016

Murayama Y, Uhlmann F (2014) Biochemical reconstitution of topological DNA binding by the cohesin ring. Nature 505: 367–371

Onn I, Aono N, Hirano M, Hirano T (2007) Reconstitution and subunit geometry of human condensin complexes. EMBO J 26: 1024–1034

Ono T, Fang Y, Spector DL, Hirano T (2004) Spatial and temporal regulation of Condensins I and II in mitotic chromosome assembly in human cells. Mol Biol Cell 15: 3296–3308

Ono T, Losada A, Hirano M, Myers MP, Neuwald AF, Hirano T (2003) Differential contributions of condensin I and condensin II to mitotic chromosome architecture in vertebrate cells. Cell 115: 109–121

Paulson JR, Hudson DF, Cisneros-Soberanis F, Earnshaw WC (2021) Mitotic chromosomes. Semin Cell Dev Biol 117: 7–29

Piazza I, Rutkowska A, Ori A, Walczak M, Metz J, Pelechano V, Beck M, Haering CH (2014) Association of condensin with chromosomes depends on DNA binding by its HEAT-repeat subunits. Nat Struct Mol Biol 21: 560–568

Sakata R, Niwa K, Ugarte La Torre D, Gu C, Tahara E, Takada S, Nishiyama T (2021) Opening of cohesin’s SMC ring is essential for timely DNA replication and DNA loop formation. Cell Rep 35: 108999

Shintomi K, Hirano T (2011) The relative ratio of condensin I to II determines chromosome shapes. Genes Dev 25: 1464–1469

Shintomi K, Hirano T (2018) Reconstitution of Mitotic Chromatids In Vitro. Curr Protoc Cell Biol 79: e48

Shintomi K, Inoue F, Watanabe H, Ohsumi K, Ohsugi M, Hirano T (2017) Mitotic chromosome assembly despite nucleosome depletion in. Science 356: 1284–1287

Shintomi K, Takahashi TS, Hirano T (2015) Reconstitution of mitotic chromatids with a minimum set of purified factors. Nat Cell Biol 17: 1014–1023

St-Pierre J, Douziech M, Bazile F, Pascariu M, Bonneil E, Sauvé V, Ratsima H, D’Amours D (2009) Polo kinase regulates mitotic chromosome condensation by hyperactivation of condensin DNA supercoiling activity. Mol Cell 34: 416–426

Suzuki K, Sako K, Akiyama K, Isoda M, Senoo C, Nakajo N, Sagata N (2015) Identification of non-Ser/Thr-Pro consensus motifs for Cdk1 and their roles in mitotic regulation of C2H2 zinc finger proteins and Ect2. Sci Rep 5: 7929

Takemoto A, Murayama A, Katano M, Urano T, Furukawa K, Yokoyama S, Yanagisawa J, Hanaoka F, Kimura K (2007) Analysis of the role of Aurora B on the chromosomal targeting of condensin I. Nucleic Acids Res 35: 2403–2412

Uhlmann F (2016) SMC complexes: from DNA to chromosomes. Nat Rev Mol Cell Biol 17: 399–412

Yoshida MM, Kinoshita K, Aizawa Y, Tane S, Yamashita D, Shintomi K, Hirano T (2022) Molecular dissection of condensin II-mediated chromosome assembly using in vitro assays. Elife [in press]

Örd M, Möll K, Agerova A, Kivi R, Faustova I, Venta R, Valk E, Loog M (2019) Multisite phosphorylation code of CDK. Nat Struct Mol Biol 26: 649–658

